# *miR*-132 controls pancreatic beta cell proliferation and survival in mouse model through the *Pten/Akt/Foxo3* signaling

**DOI:** 10.1101/233098

**Authors:** Hassan Mziaut, Georg Henniger, Katharina Ganss, Sebastian Hempel, Steffen Wolk, Johanna McChord?, Kamal Chowdhury, Klaus-Peter Knoch, Jürgen Weitz, Christian Pilarsky, Michele Solimena, Stephan Kersting

## Abstract

**Aim and hypothesis:** microRNAs (miRNAs) play an integral role in maintaining beta cell function and identity. Deciphering their targets and precise role, however, remains a challenge. In this study we aimed to identify miRNAs and their downstream targets involved in regeneration of islet beta cells following partial pancreatectomy in mice.

**Methods:** RNA from laser capture microdissected (LCM) islets of partially pancreatectomized and sham-operated mice were profiled with microarrays to identify putative miRNAs implicated in control of beta cell regeneration. Altered expression of selected miRNAs, including *miR-132*, was verified by RT-PCR. Potential targets of *miR-132* were seleced through bioinformatic data mining. Predicted *miR-132* targets were validated for their changed RNA and protein expression levels and signaling upon *miR-132* knockdown or overexpression in MIN6 cells. The ability of *miR-132* to foster beta cell proliferation *in vivo* was further assessed in pancreatectomized *miR-132*^-/-^ and control mice.

**Results:** Partial pancreatectomy significantly increased the number of BrdU+/insulin+ positive islet cells. Microarray profiling revealed 14 miRNAs, including *miR-132* and *-141*, to be significantly upregulated in LCM islets of partially pancreatectomized compared to LCM islets of control mice. In the same comparison *miR-760* was the only miRNA found to be downregulated. Changed expression of these miRNAs in islets of partially pancreatectomized mice was confirmed by RT-PCR only in the case of *miR-132* and *-141*. Based on previous knowledge of its function, we chose to focus our attention on *miR-132*. Downregulation of *miR-132* in MIN6 cells reduced proliferation while enhancing the expression of proapoptic genes, which was instead reduced in *miR-132* overexpression MIN6 cells. Microarray profiling, RT-PCR and immunoblotting of *miR-132* overexpressing MIN6 cells revealed their downregulated expression of Pten, with concomitant increased levels of pro-proliferative factors phospho-Akt and phospho-Creb as well as inactivation of pro-apoptotic Foxo3 via its phosphorylation. Finally, we show that regeneration of beta cells following partial pancreatectomy was reduced in *miR-132^-/-^* mice compared to control mice.

**Conclusions/Interpretations:** Our study provides compelling evidence for upregulation of *miR-132* being critical for regeneration of mouse islet beta cells in vivo through downregulation of its target Pten. Hence, the *miR-132*/Pten/Akt/Foxo3 signaling pathway may represent a suitable target to enhance beta cell mass.

**Research in Context:** **What is already known?**

- Several miRNAs, including *miR-132*, are known to regulate beta cell function and mass in several mouse models of diabetes *db/db, ob/ob* and high fat-diet.

**What is the key question?**

- Which are miRNAs implicated in control of beta cell regeneration upon partial pancreatectomy and how?

**What are the new findings?**

- *miR-132* is critical to promote regeneration of mouse beta cells in vivo following partial pancreatectomy
- In vitro studies in mouse MIN6 cells indicate that *miR-132* fosters beta cell proliferation by down-regulating the expression of phosphatase Pten, thereby tilting the balance between anti-apoptotic factor Akt and pro-apoptotic factor Foxo3 activities towards proliferation through regulation of their phosphorylation.

**How might this impact on clinical practice in the foreseeable future?**

- These findings strengthen the rationale for targeting the expression of *miR-132* to increase beta cell mass *in vivo* (type 2 diabetes) or *ex-vivo* (islet transplantation in type 1 diabetes) for the treatment of diabetes.

## Introduction

miRNAs belong to the class of short non-coding RNAs that regulate gene expression by annealing to 3’ untranslated region sequences in target mRNAs and inducing their post-transcriptional repression. The functional importance of miRNAs has been extensively investigated in recent years and their altered expression has been implicated in a wide range of diseases, including cancer [1], cardiovascular disease [2, 3] and diabetes [4]. In pancreatic beta cells, changed expression of miRNAs correlates with profound impairment of glucose metabolism [5] and processing of miRNAs appears to be altered in obesity, diabetes, and aging. Several miRNAs such as *miR-7*, *-21*, *-29*, *-34a*, *-132/-212*, *-184*, *-200* and *-375* have been found to be relevant for beta cell function [6]. RNA sequencing of human islets detected 346 miRNAs, including 40 which were enriched in comparison to other tissues [7]. *miR-375* is the most highly expressed miRNA in human and mouse pancreatic islets. Its down-regulation inhibits pancreatic islet development in *Xenopus laevis* [8], while its global inactivation in mice leads to decreased beta cell mass and ultimately diabetes [9, 10]. *miR-132* also plays a key role in beta cell function. Its expression is dysregulated in different mouse models of type 2 diabetes (T2D) [11–14], with its overexpression being correlated with improved glucose-stimulated insulin release from dissociated rat islet cells [13] as well as with enhanced beta cell proliferation and survival [12–14]. In primary PC12 cells, another endocrine cell model, *miR-132* controls cell survival by direct regulation of Pten, Foxo3a and p300 signaling. However, in pancreatic islets, the functional relevance of *miR-132 in vivo* and its downstream targets remain unknown. To identify the major miRNAs as well as their downstream targets involved in beta cell proliferation, we analyzed the profile of miRNAs differentially expressed after a partial pancreatectomy in comparison to sham-operated mice. We report here that the expression of 14 miRNAs were markedly increased and one miRNA was down-regulated in islets of partially pancreatectomized mice, in which beta cell proliferation was increased. Moreover, we show that down-regulation of *miR-132* expression in insulinoma MIN6 cells correlated with decreased proliferation and increased apoptosis by controlling the expression of *Pten* and its downstream effectors Akt and Foxo3 while its up-regulation had opposite effects, Finally we demonstrate that beta cell

## Methods

### Cloning of *miR-132* in *pacAd5* shuttle vectors

To produce the adenovirus overexpressing *miR-132* we used the RAPAd^®^ miRNA Adenoviral Expression System (Cellbiolabs, San Diego, CA, USA). Following the Kit’s instructions, the mmu-*miR-132* precursor sequence, obtained from www.miRBase.org (*GGGCAACCGTGGCTTTCGATTGTTACTGTGGGAACCGGAGGTAACAGTCTACAGCCATGGTCGCCC*), was PCR-amplified from genomic mouse DNA including a ca. 100bp flanking region on each side (Forward: *5’-TCGAGGATCCTCCCTGTGGGTTGCGGTGGG-3’*; Reverse: *5’-TCGAGCTAGCACATCG AATGTTGCGTCGCCGC-3’*) and cloned into the human β-globin Intron of the Kit’s *pacAd5-miR-GFP-Puro* vector via BamHI/NheI (NEB, Ipswich, MA, USA) digestion. This human β-globin intron, containing the mmu-*miR-132* precursor, was then subcloned into the Kit’s *pacAd5-CMV-eGFP* vector via PCR amplification (Forward: *5’-TGCAACCGGTGCCAG AACACAGGTACACATAT-3’*; Reverse: *5’-TGCAACCGGTCGTGCTTTGCCAAAGTGATG-3’*) and AgeI (NEB, Ipswich, MA, USA) digestion to obtain a *miR-132* overexpressing shuttle vector with a CMV promoter. The empty *pacAd5-CMV-eGFP* vector was used for the production of a control virus.

### Cell culture

Mouse MIN6 cells were kind gifts from Dr. Jun-ichi Miyazaki (Osaka University, Japan), and were grown as previously described (Miyazaki et al., 1990). MIN6 cells were cultured in 25 mmol/L glucose Dulbecco’s modified Eagle’s medium (DMEM, high glucose, GlutaMAX(TM), pyruvate) (Gibco, Thermo Fisher Scientific, Waltham, MA, USA), supplemented with 15% fetal bovine serum (Gibco, Thermo Fisher Scientific, Waltham, MA, USA), 100 U/mL penicillin, 100 U/mL streptomycin (Sigma-Aldrich, St. Louis, MO, USA) and 70μM β-Mercaptoethanol (Sigma-Aldrich, St. Louis, MO, USA), and were incubated at 37°C in a humified atmosphere containing 95% air and 5% CO_2_. HEK293T cells used to propagate the adenovirus were cultured in 25 mmol/L glucose Dulbecco’s modified Eagle’s medium (DMEM, high glucose, GlutaMAX(TM), pyruvate) (Gibco), supplemented with 10% fetal bovine serum (Gibco), 100 U/mL penicillin, 100 U/mL streptomycin (Sigma-Aldrich) and 0.1 mM non-essential amino acids (Gibco), and were incubated at 37°C in a humified atmosphere containing 95% air and 5% CO_2_.

### Adenovirus production in HEK293T Cells

The adenoviral backbone vector *pacAd5-9.2-100* from the RAPAd miRNA Adenoviral Expression System (Cellbiolabs, San Diego, CA, USA) and both shuttle vectors *pacAd5-CMV-eGFP* and *pacAd5-CMV-mir132-eGFP* were PacI linearized, and co-transfected into HEK293T cells using the X-tremeGENE 9 DNA Transfection Reagent (Roche, Basel, Switzerland) and amplified, until a sufficient amount of virus supernatant was available for experiments, which was then aliquoted and stored at −80°C. Virus titers were determined with the Adeno-X^TM^ Rapid Titer Kit (Clontech, Mountain View, CA, USA) All steps were carried out according to the manufacturer’s instructions.

### Altered *miR-132* expression with siRNAs or with adenovirus

The single-stranded RNA used in to silence *miR-132* consisted of 24-nucleotide length oligo: *5’-CGACCAUGGCUGUAGACUGUUACC-3’* and as control, we used a scrambled oligonucleotide. MIN6 cells were seeded in a 6-well plate as described above on day 1. For the silencing of *miR-132*, cells were transfected with 100 nM of anti-*miR-132* or control siRNAs using Dharmafect4 as transfection reagent on day 2. For the overexpression of *miR-132*, MIN6 cells were transduced with an adenovirus in which *miR-132* was driven by the CMV promoter at MOI of 6,400. On day 4 post RNA silencing or adenoviral transduction, cells were harvested proceded for BrdU labeling or western blotting as described below.

### Cell extraction and immunoblotting

MIN6 cells were harvested at 4°C in RIPA buffer [50 mM Tris⋅HCl, pH 8.0, 150 mM NaCl, 1% Nonidet P-40, 0.1% SDS, 0.5% sodium deoxycholate, and protease inhibitor mixture (Sigma)] for total protein extraction. Insoluble material was removed by centrifugation. Aliquots of 20 ug were separated by SDS PAGE, as described (Mziaut et al., 2008). The source, species and dilutions of antibodies used immunoblotting are described in ESM Table 6.

### Mouse studies

*miR-132/-212^-/-^* mice were backcrossed into the *C57Bl/6N* background for at least seven generations, as previously described (Ucar et al., 2010). Mutant and wildtype male *C57Bl/6N* mice with an age of 13-19 weeks and a body weight of 28-34 g were used for the experiments. All animal protocols were approved by the institutional animal care and use committee and all experiments were performed in accordance with relevant guidelines and regulations.

Mice underwent general anesthesia by mask inhalation of isoflurane using a small rodents’ anesthesia unit (Harvard Apparatus Ltd., Holliston, MA, USA) with an isoflurane (Baxter Deutschland GmbH, Unterschleißheim) concentration of 4.5–5% for induction and 2-2.5% for maintenance of anesthesia with an airflow rate of 200 ml/min. Perioperative analgesia was accomplished using buprenorphin (0.05 mg/kg bodyweight) administered subcutaneously. The abdomen was opened through an upper midline incision. The spleen and the entire splenic portion of the pancreas were surgically removed while the mesenteric pancreas between the portal vein and the duodenum was left intact. The remnant was defined as the pancreatic tissue within 1-2 mm of the common bile duct that extends from the duct to the first portion of the duodenum. This remnant is the upper portion of the head of the pancreas. This procedure results in approximately 75% pancreatectomy, as confirmed by weighing the removed and remnant portions of the pancreas (Mziaut et al., 2008). Sham operations were performed by removing the spleen while leaving the pancreas intact. At the end of surgery, Alzet 1007D miniosmotic pumps (Alzet^®^, Cupertino, CA, USA) were implanted i.p. to deliver 50 µg⋅µl^-1^ BrdU (Sigma, St. Louis, MO, USA) in 50% DMSO at a rate of 0.5 µl⋅h^-1^ for 7 days. Blood glucose levels were measured daily from the tail vein with a Glucotrend glucometer (Roche Diagnostics, Basel, Switzerland).

### Intraperitoneal glucose tolerance test (IpGTT)

IpGTT were performed two days before surgery and six days after surgery to assess differences between wildtype and mutant mice and between pancreatectomized and sham-operated animals. After 10 hours overnight fast, mice were injected with 2.0 g/kg body weight of 20% glucose solution. Blood glucose levels were measured from the tail vein at 0, 15, 30, 60, 120 and 180 minutes after glucose injection.

The pancreatic remnants and the sham-operated pancreata were harvested 7 days post-surgery. Mice were anesthetized with isoflurane as described above and the abdominal incision was re-opened. After fixation by intracardial perfusion with 4% paraformaldehyde, mouse pancreata were removed, further fixed overnight in 10% neutral formalin, and embedded in paraffin. Sections were cut at 5 μm and BrdU staining in MIN6 Cells was performed as described in (Mziaut et al., 2008).

### Real Time-PCR

cDNA samples were obtained by reverse transcription of 1ug total RNA using the M-MLV Reverse Transcriptase (Promega, USA, WI, Madison). Quantitative real time-PCR was then performed with the GoTaq® qPCR Master Mix (Promega, USA, WI, Madison) according to the manufacturers instruction using the oligonucletides listed in ESM Table 7.

### Transcriptomic profiling of mouse islets

Total RNA was isolated from the islets of 16 - 19 -week-old wild-type and *miR-132^-/-^* mice (6 mice/group) using RNeasy (Qiagen, Hilden, Germany). For microarray analysis, 100 ng of islet RNA was amplified with the Illumina® Total Prep RNA Amplification Kit (Ambion, Inc., Austin, TX, USA) and cRNA was labeled with biotin-UTP as previously described (Mziaut et al., 2016).

### IMicroarray analysis and data mining

MicroArray data processing and quality control was done with GeneSpring GX 13. After quantile normalization significantly differentially expressed genes were obtained by volcano plot filtering for FC≥|1.5| and p<0.05. Functional analysis was done using Ingenuity Pathway Analysis (IPA, Qiagen, Hilden, Germany).

### Statistical analysis

Statistical analyses were performed by using the unpaired Student’s *t* test unless otherwise stated. Results are presented as mean SE unless otherwise stated. A value of *P* < 0.05 was considered significant. Error bars show standard deviations from at least three independent experiments unless otherwise stated. Histograms were prepared with Microsoft Excel (Microsoft, Redmont, WA, USA) or GraphPad Prism.

## Results

### *miR-132* is upregulated in proliferating beta cells

To identify key miRNAs involved in beta cell proliferation, we used partially pancreatectomized mice (n=3) as a model for beta cell regeneration (Fig. 1A). The removal of 70-80% of the pancreas is a well-established procedure for inducing the replication of beta cells in the remaining pancreas [15, 16]. Proliferating cells were stained with 25µg/hr BrdU continuously delivered by an osmotic mini-pump implanted in the abdomen at the time of pancreatectomy. This approach ensures the labeling of every dividing cell [16]. As controls, 4 mice were similarly implanted with an osmotic mini-pump for BrdU delivery, but only underwent total splenectomy. Seven days post-surgery, all mice were sacrificed, their pancreas excised for serial sectioning (40 sections/mouse) and staining with cresyl-violet to locate the islets. Adjacent, unstained sections were then used to count BrdU^+^ cells in the islet cores, which in rodents consists mainly of beta cells [15] (Fig. 1B and C) prior to islet core excision by laser capture microdissection (LCM) (Fig. 1 A and B). As expected, in partially pancreatectomized mice the fraction of islet core BrdU^+^ cells/total islet core cells, as determined by nuclear counting, was significantly higher than in sham-operated mice (Fig. 1C). RNA extracted from LCM islet cores was then profiled using microarrays. Fourteen miRNAs were found to be differently expressed (cut-off values: p= 0.05; FC≥1.5) in the islet cores of partially pancreatectomized mice compared to sham operated mice (Table 1). All these miRNAs were up-regulated, except *miR-760*, which was reduced by 2.28 fold. Expression levels of all 14 differentially expressed miRNAs were further quantified by real-time PCR (RT-PCR). With this analysis, only the significant changed expression of *miR-132* and *miR-141* could be validated (ESM Table 1). Given the role of *miR-132* in the cell replication *in vitro* of primary islet cells [13], and other cell types, including glioma cells [17] and epidermal keratinocytes [18], we focus our attention on the potential involvement and mode of action of miR in the regulation of beta cell proliferation.

**Figure 1.**
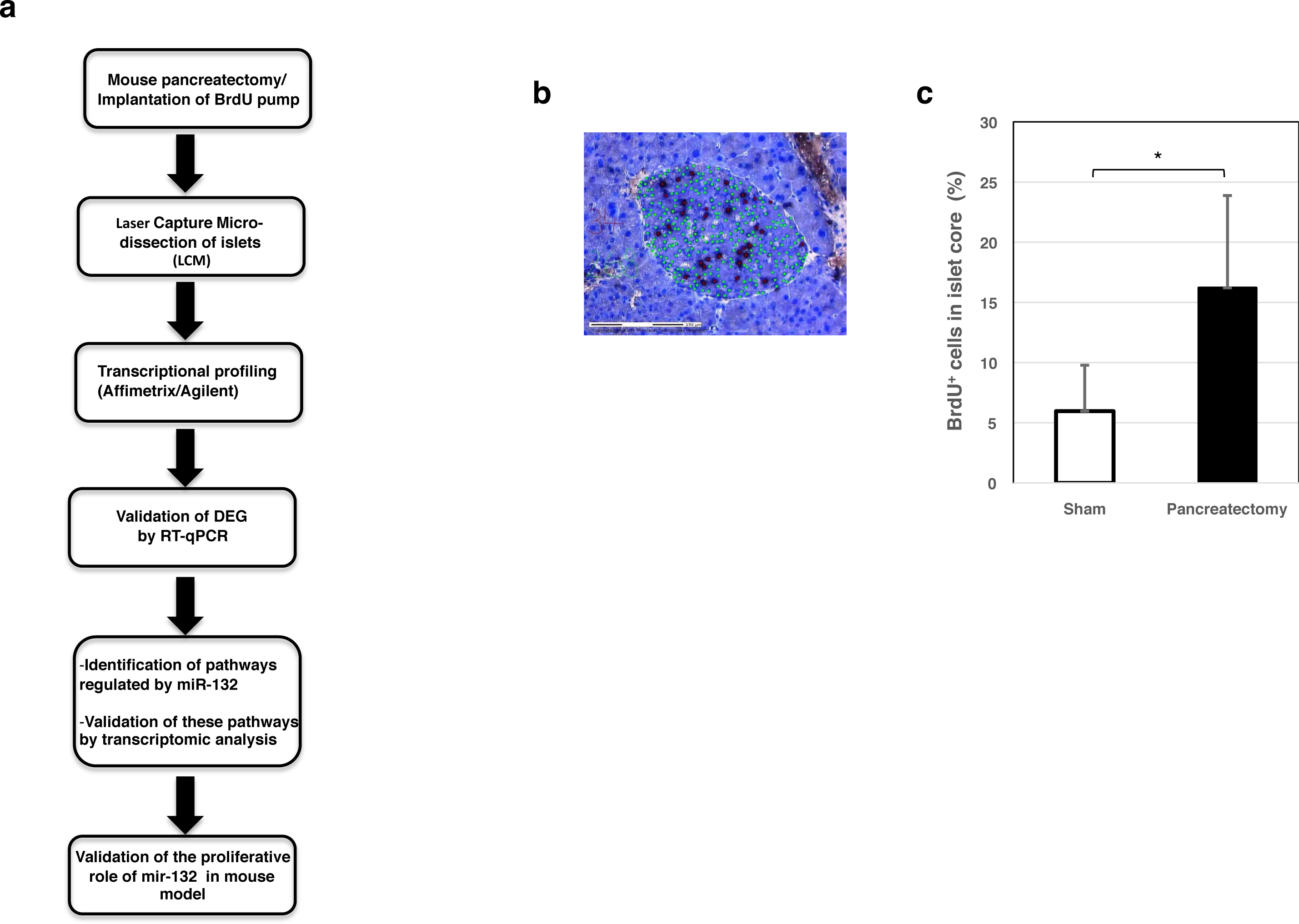
*miR-132* expression is induced in pancreatectomized mice. Overview of the experimental design (A). Black6 mice were pancreatectomized and pump was inserted in the abdomen for continous BrdU administration over a week. Pancreatic sections were stained with cresyl-violet dye to reveal the islet core and labeled with DAPI (pseudogreen) and with anti-BrdU antibody (pseudoblack) (B top panel). The islets core was excised by laser capture micro-dissection (LCM) and processed for micro-RNAs isolation and profiling (B bottom panel). (C) Percentage of BrdU^+^ cells in islet cores. Experiment included 7 mice of which 4 were shamp operated and 3 were pancreatectomized. 40 slices/group were counted for BrdU^+^ cells. *, *P* < 0.05.

**Table 1:** miRNAs differentially expressed in islet cores of partially pancreatectomized mice ranked by fold change (absolute value)

| Systematic name | p-value | FC | Px1 | Px1 | Px3 | S1 | S2 | S3 | S4 |
| --- | --- | --- | --- | --- | --- | --- | --- | --- | --- |
| mmu-mir-205 | 0.009 | 4.91 | 6.5011 | 12.7521 | 9.4035 | 1 | 5.34742 | 2.0243 | 1 |
| mmu-mir-132 | 0.005 | 4.63 | 85.442 | 90.0032 | 71.439 | 26.371 | 16.9079 | 22.282 | 8.6324 |
| mmu-mir-186 | 0.016 | 2.5 | 4.0622 | 5.34086 | 3.1434 | 1.7227 | 1.1133 | 1.9388 | 1.686 |
| mmu-mir-760 | 0.041 | - 2.28 | 12.088 | 13.0156 | 4.6472 | 12.305 | 37.794 | 19.394 | 17.248 |
| mmu-mir-129-3p | 0.022 | 1.99 | 213.55 | 272.406 | 121.26 | 118.28 | 100.958 | 92.346 | 68.376 |
| mmu-mir-690 | 0.03 | 1.79 | 49.413 | 48.5373 | 30.88 | 23.957 | 39.5096 | 16.126 | 17.206 |
| mmu-mir-130b | 0.014 | 1.77 | 42.937 | 40.5664 | 29.545 | 22.405 | 28.2766 | 18.247 | 14.679 |
| mmu-mir-300* | 0.045 | 1.77 | 2.9997 | 4.06448 | 2.006 | 1.8972 | 1.78003 | 1.8814 | 1 |
| mmu-mir-598 | 0.049 | 1.75 | 18.357 | 22.9878 | 14.596 | 12.314 | 10.6534 | 12.703 | 6.3097 |
| mmu-mir-431 | 0.013 | 1.65 | 23.393 | 27.3673 | 17.432 | 14.414 | 15.546 | 13.637 | 9.7251 |
| mmu-mir-329 | 0.035 | 1.64 | 25.756 | 29.392 | 19.037 | 17.004 | 16.0322 | 15.797 | 9.9605 |
| mmu-mir-154* | 0.029 | 1.63 | 19.404 | 19.675 | 15.306 | 10.545 | 11.4188 | 13.067 | 8.2187 |
| mmu-mir-141 | 0.04 | 1.59 | 878.04 | 970.708 | 655.42 | 566.11 | 672.314 | 515.55 | 318.15 |
| mmu-mir-200c | 0.039 | 1.58 | 976.04 | 997.919 | 672.69 | 608.97 | 681.16 | 530.91 | 361.34 |
| mmu-mir-375 | 0.107 | 1.48 | 6291.99 | 7241.707 | 4643.13 | 4778.29 | 4398.79 | 4440.34 | 2461.28 |

### *miR-132* affects proliferation and survival of insulinoma MIN6 cells

At first, we down-regulated *miR-132* expression in mouse insulinoma MIN6 cells using an anti-microRNA approach [19]. Reduced expression of *miR-132* by 90% (n=3, Fig. 2A), as assessed by RT-PCR, correlated with a slight, but significant reduction in the percentage of BrdU^+^ MIN6 cells in comparison to cells [n=6, 24.48±1.78 % (6,088 BrdU^+^ out of 25,246 cells) vs. 27.35±2.23% (7,271 BrdU^+^ cells out of 27,413), p-value= 0.039)], which had been transfected with a control oligonucleotide (Fig. 2B and C). *miR-132* depletion correlated also with increased detection of cleaved Caspase-9 (n= 6; Fig. 2D and E), while the levels of cleaved Caspase-3 were not significantly changed. Conversely, overexpression of *miR-132* with a bi-cistronic adenovirus vector encoding also for eGFP (n= 3 Fig. 2F) was not associated with a further increase in the proliferation of MIN6 cells, presumably due to their neoplastic state (n= 3, Fig. 2G and H). However, overexpression of *miR-132* correlated with reduced levels of pro- and cleaved Caspase-9 (n= 6, Fig. 2I and J), consistent with *miR-132* being anti-apoptotic.

**Figure 2.**
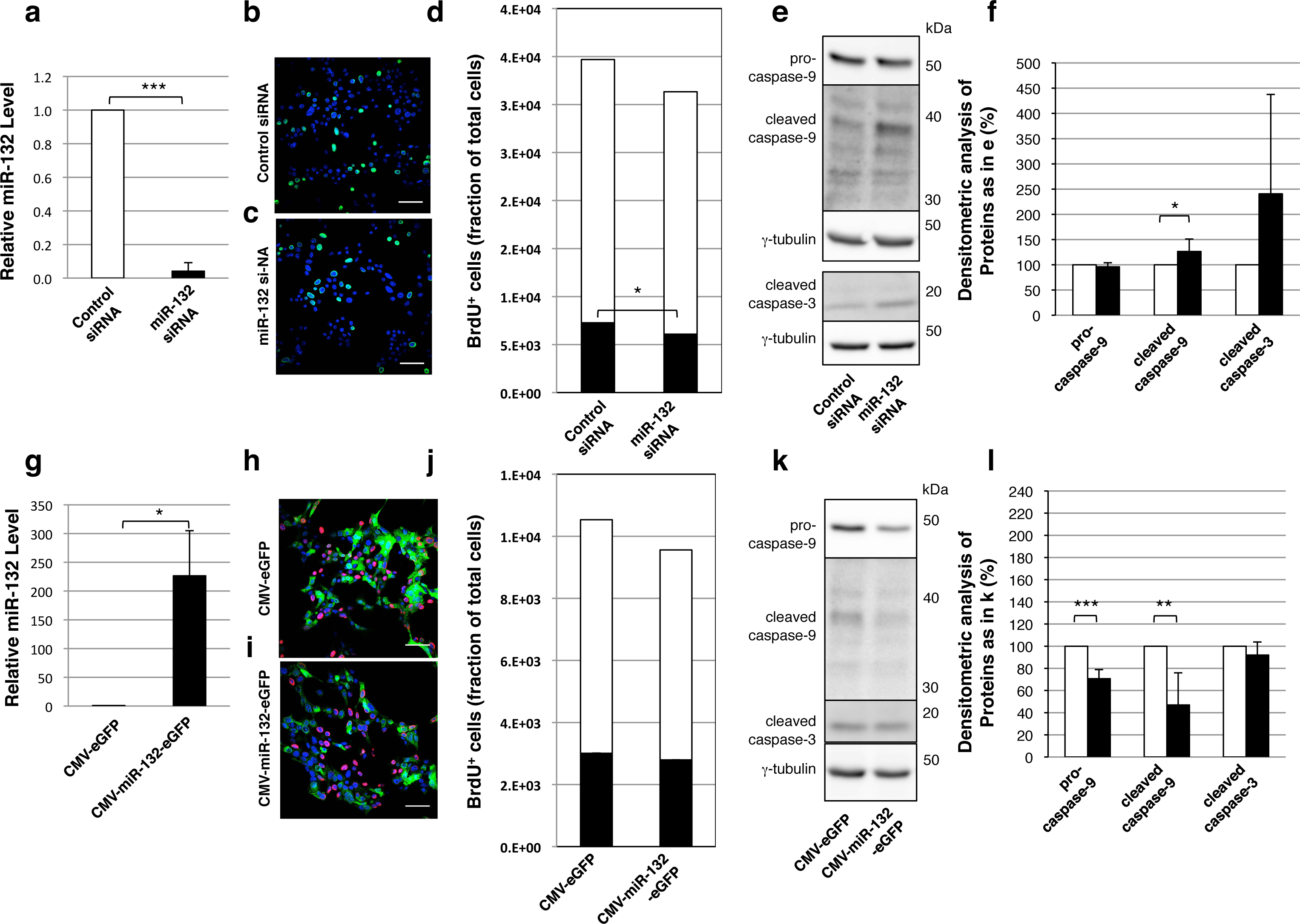
*miR-132* regulates the proliferation and cell survival in MIN6 cells. Down-regulation of *miR-132* with siRNA (A). Analysis of the proliferation (B, C) and apoptosis (D, E) of *miR-132* depleted cells respectively by quantification of BrdU positive cells and immunoreactivity against cleaved caspase 3 and 9. Adenovirus mediated expression of *miR-132* in MIN6 cells (F) and its role on the proliferation (G, H) and apoptosis (I, J) of MIN6 cells. Experiments were performed 3 times for (A and F) and 6 times for (D-E and I-J). *, *p* < 0.05, **<0.01, **<0.001. Scale bar = 50µm

### *miR-132* regulates the expression of *Pten* and *Mapk1*

Next, we aimed to uncover down-stream targets of *miR-132* in MIN6 cells. Microarray gene expression analysis of *miR-132* overexpressing MIN6 cells identified 345 unique differentially expressed genes (cut-off values: p<0.05, FC ≥1.5), with 194 (56.2%) being down regulated and 151 (43.8%) up-regulated (Fig. 3A and ESM Table 2). Querying the TargetScan Mouse 7.1 database 35 of the down- and 2 of the up-regulated genes were predicted to contain highly conserved binding sites for *miR-132* (Fig. 3A and ESM Table 3). Further analysis of regulated genes with Ingenuity Pathway Analysis revealed 8 regulated pathways (Fig. 3B and ESM Table 4), which included 26 of the 345 differentially expressed genes (Fig. 3C). Among these 26 differentially expressed genes, the top 10 most represented genes in the 8 signaling pathways were selected for further validation of their mRNA levels by RT-PCR (ESM Table 5). As control, we also assessed the mRNA expression levels of *Rasa1*, an established target of *miR-132* (Anand et al., 2010). As shown in Fig. 3D, 6 out of 10 of the selected genes, namely *Mapk1/Erk2*, *Pten*, *Nras*, *Pik3r1*, *Gnb1* and *Gnb5* were confirmed to be downregulated upon *miR-132* overexpression. Three of them, *Mapk1, Pten* and *Gnb1,* were also among the 37 predicted targets for *miR-132* binding (ESM Table 6). Notably, *Mapk1,* also known as *Erk2,* is a serine-threonine kinase located downstream of the tumor-suppressor phosphatase *Pten* and both genes play a critical role in control of cell proliferation and survival (Deb et al., 2014; Ya-Chun et al., 2015).

**Figure 3.**
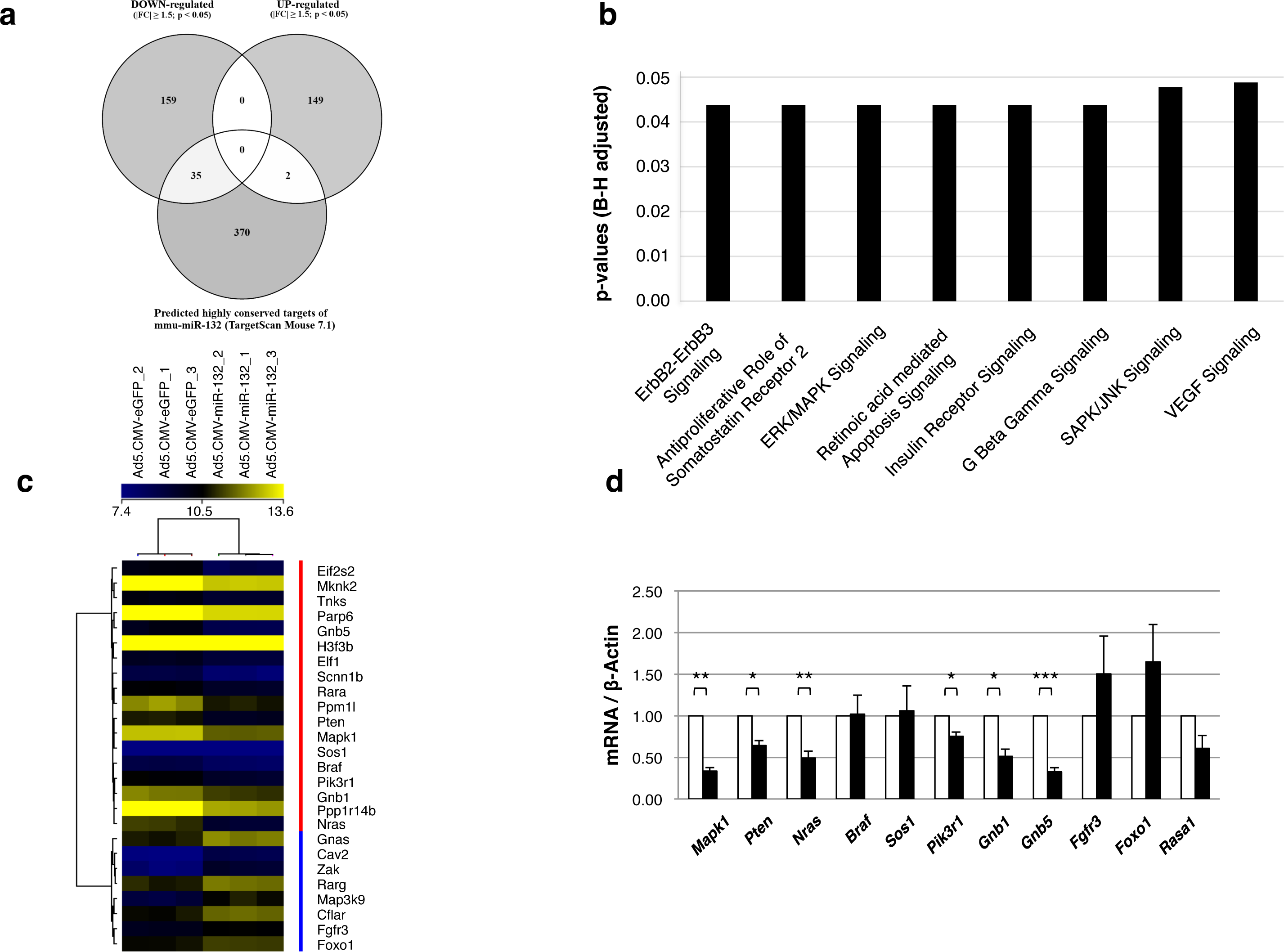
Identification and validation of *miR-132* target genes in MIN6 cells. Venn diagram showing the number of differentially expressed (DE) genes in MIN6 cells overexpressing *miR-132* (A). Ingenuity Pathway Analysis of significantly altered pathways in cells expressing *miR-132* (B). Top ranked DE genes with known role in cell proliferation and survival (C). Validation by RT-PCR of selected DE genes uncovered by microarray analysis (D).

### miR-132 regulates Pten signaling in MIN6 cells

Next, we tested whether overexpression of *miR-132* affected the protein levels of the Pten and Mapk1/Erk2. Immunoblotting of MIN6 cells transduced with the *miR-132*/*eGFP* viral vector confirmed the down-regulation of Pten in parallel with up-regulation of its targets Akt and phospho-Akt (S473) (Fig. 4A and B), while mRNA levels of Akt were unchanged (ESM Fig. 1). Furthermore, levels of the Akt1 substrate Creb and phospho-Creb (S133) were unchanged, but the phospho-Creb/Creb ratio was also increased. Likewise, overexpression of *miR-132* correlated with reduced expression of Mapk1/Erk2, phospho-Mapk1/Erk2 and Rasa1/RasGAP, but not of Erk1 and phospho-Erk1 (Fig. 4C and D).

**Figure 4.**
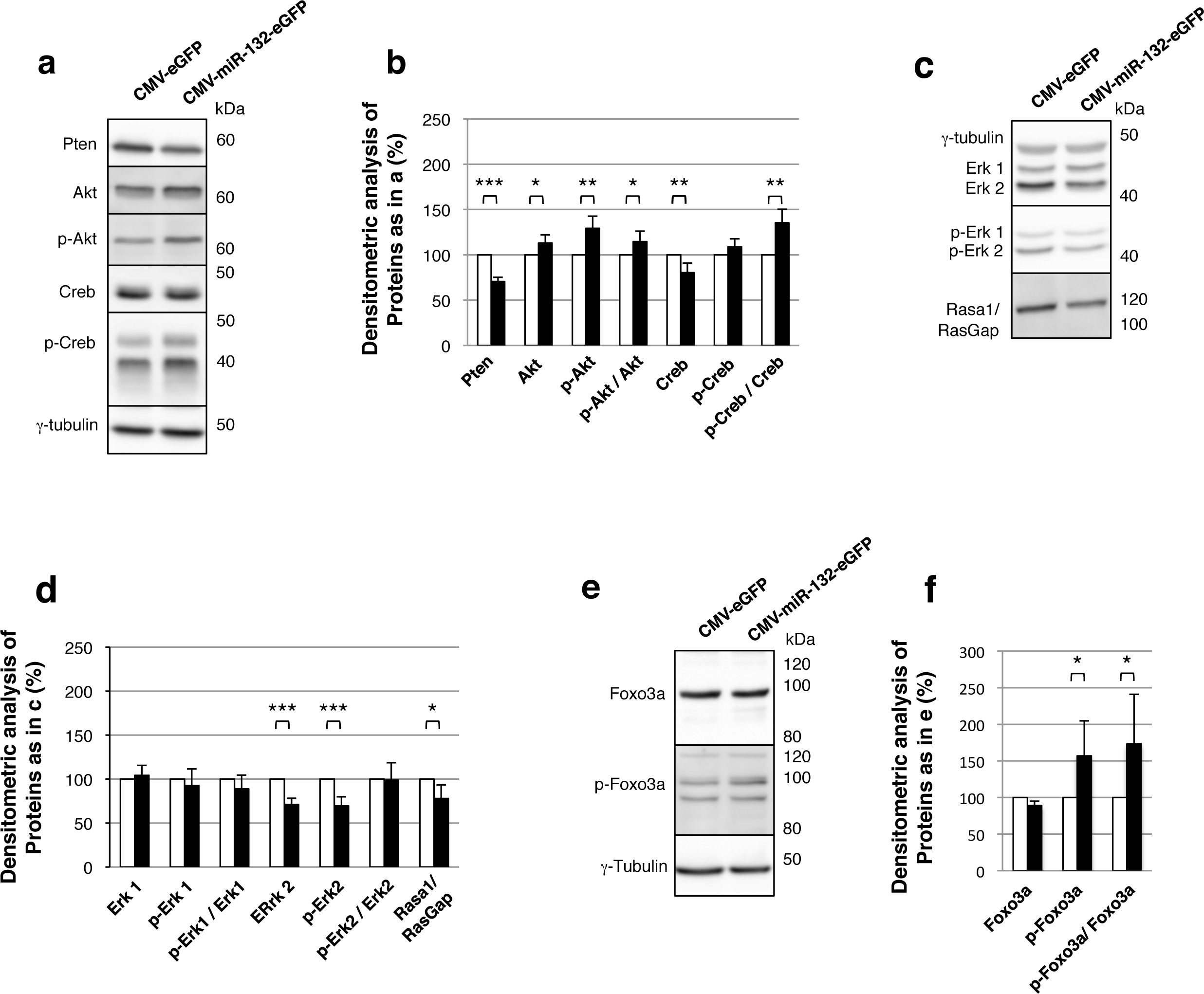
Validation of gene profiling data in MIN6 Cells overexpressing miR132. Immunoblotting with the indicated antibodies on cell extracts from Min6 cells transduced with a adenovirus vector plasmid carrying a CMV promoter driving the expression of control or *miR-132* in tandem with the reporter protein GFP (A, C and E). (B, D and F) Densitometric quantification of the indicated proteins as shown in A and C and E. Immunoblotting and quantification were performed from 6 experiments.

As down-regulation of *miR-132* correlated with elevated cleaved Caspase-9 levels, we further tested whether its overexpression affected the expression of *Foxo3a,* a key mediator of apoptosis. Immunoblotting for Foxo3a showed that its phosphorylation, which inhibits its activity,was increased (Fig. 4E-F), although the overall levels of its mRNA (ESM Fig. 1) and protein (Fig. 4E-F) were not changed.

### Impaired beta cell proliferation in *miR-132* knock-out mice

Finally, to verify that *miR-132* positively affects beta cell regeneration *in vivo*, we investigated beta cell proliferation in partially pancreatectomized or sham operated *miR-132*^-/-^ mice and control littermates (6 mice/group), as described previously (Mziaut et al., 2008). Intraperitoneal glucose tolerance test prior and 6 days after surgery showed no difference between control and *miR-132^-^*^/-^ mice (Fig. 5A and B). Daily blood glucose measurements, in particular, showed a comparable slight decrease of glycaemia in partially pancreatectomized wild-type and *miR-132*^-/-^ mice relative to sham operated mice in the first day post-surgery, followed by a complete normalization of glycaemia by the end of the 1-week-long protocol (ESM Fig. 2). Seven days after surgery, the mice were sacrificed, the remnant pancreas excised, and BrdU^+^/insulin^+^ beta cells were counted (Fig. 5C-F and Table 2). As assessed by immunostaining for insulin, the average number of beta cells/islet in wt (31,9 beta cells/islet) and *miR-132*^-/-^ (31,7 beta cells/islet) was increased in partially pancreatectomized mice relative to their sham-operated counterparts (wt: 23,8 beta cells/islet; *miR-132*^-/-^: 26,8 beta cells/islet). Likewise, the number BrdU^+^ insulin^+^ beta cells was increased in both groups of partially pancreatectomized mice compared to sham operated mice (Fig. 5G, Table 2). However, in partially pancreatectomized *miR-132*^-/-^ mice there were fewer BrdU^+^/insulin^+^ cells than in partially pancreatectomized wt mice (Fig. 5G, Table 2). These data provide conclusive evidence for *miR-132* exerting a positive role for beta cell regeneration *in-vivo*.

**Figure 5.**
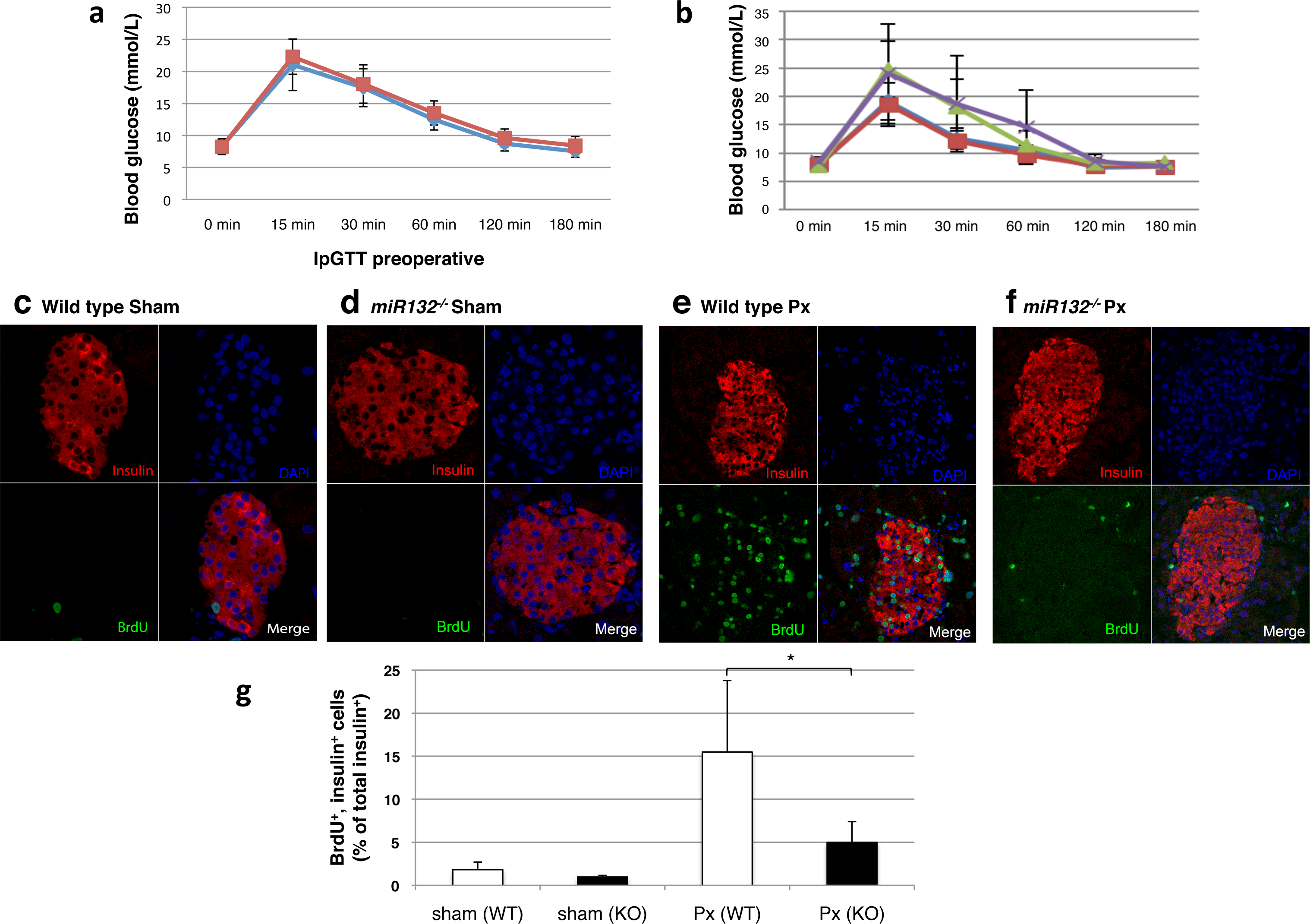
Impaired pancreatic beta cells regeneration in *miR-132*^-/-^ mice. IPGTT two days before (A) and six days after (B) surgery in wildtype and *miR-132*^-/-^ mice, either sham-operated or pancreatectomized. Immunomicroscopy on paraffin sections of the remnant pancreas from sham-operated wildtype (C) or *miR-132*^−/−^ (D) mice, and partially pancreatectomized wildtype (E) or *miR-132*^−/−^ mice (F) after one week of continuous administration of BrdU. Sections were immunolabeled with anti-insulin (pseudored) and anti-BrdU (pseudogreen) antibodies. Nuclei were counterstained with DAPI (pseudoblue). Percentage of insulin^+^ and BrdU^+^ cells (G). Three independent series with two mice per condition were used and at least 50 islets per mouse for a total of 250- 300 islets per group were counted. *, *p* < 0.05.

## Discussion

*miR-132* is known *to* control many cellular processes in various tissues including neuronal morphogenesis and regulation of circadian rhythm. *miR-132* altered expression correlated with several neurological disorders, such as Alzheimer’s and Huntington’s diseases (Wang et al., 2013; Lee et al., 2011). Thus, most of our acknowledge about *miR-132*regulation and biological functions emerged from studies performed on neural cells, while not much is already known about the downstream target of *miR-132* in pancreatic beta cells. *miR-132* and *miR-212* have identical seed sequences and potentially have targets in common. Here we have identified *miR-132* as one of the mostly up-regulated miRNAs with a 5 fold expression change in pancreatectomized mice, a condition known to induce beta cell proliferation. This finding is in agreement with previous data showing induced *miR-132* expression in different models of type 2 diabetes, including *db/db*, high-fat diet-fed (Nesca et al., 2013) and *ob/ob* mice (Zhao et al., 2009; Poy et al., 2009). Among our list of differentially expressed miRNAs in islets of partially pancreatectomized mice, *miR-205* showed the greatest change (5 fold). Interestingly, *miR-205* has also been described to be the miRNA with the highest expression change in hepatocytes of mice with obesity-induced type 2 diabetes (Zhao et al., 2009). On the other hand, our analysis did not reveal a significant change in the expression of *miR-375*, a miRNA abundant in pancreatic islets known to regulate insulin secretion and beta cell proliferation (Poy et al., 2009).

Previous work has shown that *miR-132* is highly expressed in neurons and may regulate their differentiation (Vo et al. 2005; Wayman et al. 2008). More recent work on primary neurons and PC12 cells has shown that *miR-132* controls cell survival by direct regulation of Pten, Foxo3a and p300 - all proteins involved in Alzheimer neuro-degeneration (Wong et al., 2013). To uncover the down stream target of *miR-132* in beta cells, we first investigated whether *miR-132* has a proliferative role in insulinoma MIN6 cells. We found that *miR-132* down-regulation with anti-*miR-132* correlated with slight but significant inhibition of MIN6 cells proliferation. The inhibition of the expression of *miR-132* showed also an increase in cleaved Caspase-9-mediated apoptosis. Conversely, its up-regulation has a protective effect by reducing Caspase-9 processing. However, the increase in the expression of *miR-132* did not correlate with an enhancement in the proliferative rate of MIN6 cells, presumably due to their higher proliferative fate. To uncover the downstream targets of *miR-132*, we analyzed the expression pattern of differentially expressed genes in cells over-expressing *miR-132*. *Pten* expression which is known to inversely correlate with cell survival, was significantly down-regulated in agreement with our data showing decreased apoptosis upon overexpression of *miR-132*. The reduced expression of *Pten* upon overexpression of *miR-132* was confirmed at the protein level. Importantly, reduction of Pten correlated with increased phosphorylation of Akt (P-Akt) and Foxo3. P-Akt is a major activator of Foxo family proteins, which are members of the Forkhead superfamily of winged helix transcription factors controlling cellular metabolism, stress responses, DNA damage repair and cell death. P-Foxo3 can be phosphorylated on Thr32, Ser253 and Ser315 which promotes its and association with 14-3-3 proteins for retention in the cytoplasm, and thereby inhibition of its transcriptional activity (Brunet et al., 1999).

Mapk1/Erk2 (mitogen-activated protein kinase 1) expression is reduced in cells over-expressing *miR-132* and survey of Erk2 activation reveals reduced phosphorylation of Erk2 without change in its activation state (as shown by the ratio of p-Erk2/Erk2) suggesting that the Ras/Raf/Erk1/2 is not preponderant in this biological process.

A considerable number of miRNAs have been associated with pancreatic beta cell development by affecting proliferation or differentiation (e.g., *miR-375* (Poy et al., 2009), *miR-7* (Wang et al., 2013), *miR-124a* (Baroukh et al., 2007), *miR-24* (Vijayaraghavan et al., 2014), *let-7a* (Gurung et al., 2014), *miR-26a* (Fu et al., 2013), *miR-184* (Tattikota et al., 2014), *miR-195*, *miR-15*, *miR-16* (Joglekar et al., 2007) and *miR-132* (Guay et al., 2015 and Zhao 2009). Among those miRNAs, *miR-132* was consistently differentially expressed in various T2D models in which beta cells were challenged by increased metabolic demand, condition known to promote beta cell proliferation, including obesity induced diabetes (Regazzi, Zhao et al., 2009). In mouse model, constitutive deletion of *miR-132* resulted in mice with deficient endocrine development (Ucar et al., 2010). A specific deletion of *miR-13*/*212* locus in adult hippocampus with a retrovirus expressing Cre recombinase caused a dramatic decrease in dendrite length, arborization and spine density, suggesting that *miR-132*/*212* is required for normal dendritic maturation in adult hippocampal neurons (Magill et al., 2010). Here we demonstrate that regeneration of beta cells in pancreatectomized *miR-132*^−/−^ mice is reduced, conceivably through its control of the Pten/Pi3K/Akt signaling.

In conclusion, we have discovered a miRNA that acts as sensor of external changes and controls the major signaling pathway regulating cell proliferation and survival at their node. Efforts to develop targeted therapies have not been fully successful, mainly because of extensive networking between pathway suppressors. Thus, our findings suggest that *miR-132* could be a suitable candidate for therapeutic intervention to prevent beta cell damages and death as well as in improving beta cell regeneration. Moreover, The identification of a number of targets allowed us to propose mechanisms by which *miR-132* is involved in these processes and eventually, could set up the ground toward the identification of the extracellular signal and the biological processes that favors a signaling pathway over another one in future studies.

## Acknowledgements

We are grateful to department of general thoracic and vascular surgery at the faculty of medicine in Dresden for their support.

## Data availability

The data are available on request.

## Contribution statement

HM, MS and SK conceived the study and the experimental design; GH profiled gene expression, performed the data mining and validated the target genes with the help of KG, JM, KPK and SW; SH and JM performed the pancreatectomy, implanted the BrdU pump and immunostained pancreatic islets under the supervision of SK. KC generated and provided the *miR-132*^-/-^ mice. HM, MS and SK wrote the manuscript. HM, MS and SK are responsible for the integrity of the study.

---

**Supplementary Figure 1. Validation of additional differentially expressed genes in MIN6 cells overexpressing *miR-132* by RT-PCR**.

**Supplementary Figure 2. Glycaemic profile of sham and partially pancreatectomized control or *miR-132*^-/-^ mice pre- and up to 6 days post-surgery**.

## Abbreviations

miRNA: microRNA
*miR-132*: microRNA 132
Pten: Phosphatase and Tensin homolog
*Akt*: proteine kinase B
*Foxo3*: *Forkhead box O3*;
RT-PCR: Reverse transcription polymerase chain reaction
T2D: type 2 diabetes
BrdU: bromodeoxyuridine
LCM: laser capture microscopy
IPA: Ingenuity Pathway Analysis
*Pi3k*: Phosphatidylinositol-4,5-bisphosphate 3-kinase
*Ras*: Rat sarcoma
*Raf*: rapidly accelerated fibrosarcoma
*Mapk*: mitogen-activated protein kinase
*Sos1*: Son of sevenless homolog 1
*Gnb1*: G Protein Subunit Beta 1
*Nras*: Neuroblastoma RAS viral oncogene homolog
*Pik3r1*: Phosphoinositide-3-Kinase Regulatory Subunit 1
*Gnb*: G Protein Subunit Beta
*Fgfr3*: *fibroblast growth factor receptor 3*
*Creb*: cAMP response element-binding protein

